# Exploring the link between Parkinson’s disease and Diabetes Mellitus in *Drosophila*

**DOI:** 10.1101/2022.02.18.481049

**Authors:** Francisco José Sanz, Cristina Solana-Manrique, Joaquín Lilao-Garzón, Yeray Brito-Casillas, Silvia Muñoz-Descalzo, Nuria Paricio

## Abstract

Parkinson’s disease (PD) is the second most common neurodegenerative disease. Diabetes mellitus (DM) is a metabolic disease characterized by high levels of glucose in blood. Recent epidemiological studies are highlighting the link between both diseases; it is even considered that DM might be a risk factor for PD. To further investigate the likely relation of these diseases, we have used a *Drosophila* PD model based on inactivation of the *DJ-1β* gene (ortholog of human *DJ-1*), and diet-induced *Drosophila* and mouse T2DM models, together with human neuron-like cells. T2DM models were obtained by feeding flies with a high sugar containing medium, and mice with a high fat diet. Our results showed that both fly models exhibit common phenotypes such as alterations in carbohydrate homeostasis, mitochondrial dysfunction or motor defects, among others. In addition, we demonstrated that T2DM might be a risk factor of developing PD since our diet-induced fly and mouse T2DM models present DA neurodegeneration, a hallmark of PD. We have also confirmed that neurodegeneration is caused by increased glucose levels, which has detrimental effects in human neuron-like cells by triggering apoptosis and leading to cell death. Besides, the observed phenotypes were exacerbated in *DJ-1β* mutants cultured in the high sugared medium, indicating that DJ-1 might have a role in carbohydrate homeostasis. Finally, we have confirmed that metformin, an antidiabetic drug, is a potential candidate for PD treatment and that it could prevent PD onset in T2DM model flies. This result supports antidiabetic drugs as promising PD therapeutics.

## INTRODUCTION

Parkinson’s disease (PD) is the second most common neurodegenerative disease, and the first motor disorder affecting more than 1-3% of the worldwide population over 60 years (Tysnes & Storstein, 2017; Xiong & Yu, 2018). Its major pathological hallmark is the selective loss of dopaminergic (DA) neurons in the substantia nigra pars compacta, which leads to PD-characteristic motor deficits. However, the exact cause of this neurodegeneration remains unknown (Athauda & Foltynie, 2016). To date, multiple pathways appear to contribute to PD onset and progression, like accumulation of misfolded protein aggregates, mitochondrial dysfunction, oxidative stress (OS), neuroinflammation, endoplasmic reticulum (ER) stress, or genetic mutations. In addition, metabolic alterations have been recently shown to play an important role in PD physiopathology. Specifically, a possible link between glucose metabolism alterations and neurodegeneration has been demonstrated (Anandhan et al., 2017; Maiti et al., 2017; Solana-Manrique et al., 2020, 2022; Sonninen et al., 2020). Even though the majority of PD cases are sporadic (sPD), several genes responsible for familial PD forms (fPD) have been identified (Bandres-Ciga et al., 2020). Interestingly, the study of their functions has provided useful information to decipher several molecular mechanisms underlying PD (Chai & Lim, 2013). Among them, *DJ-1*, initially described as an oncogene, is a causative gene of early-onset fPD (Bonifati et al., 2003). However, further studies demonstrated that the DJ-1 protein have additional functions such as antioxidant, via free-radical scavenging and transcriptional regulation of antioxidant genes, as mitochondrial function modulator, deglycase, redox-dependent molecular chaperone, and as a factor involved in proteolysis and metabolism (Solana-Manrique et al., 2020, 2022; Strobbe et al., 2018).

Diabetes mellitus (DM) is a metabolic disease characterized by high levels of glucose in blood, due to a diminished insulin production or to the appearance of insulin resistance. There are, mainly, two types of DM: type 1 DM (T1DM) and type 2 DM (T2DM). T1DM is due to the autoimmune loss of pancreatic β-cells, whereas T2DM is caused by the appearance of insulin resistance and a progressive decrease of insulin production in β-cells (American Diabetes Association, 2013; Hassan et al., 2020). Interestingly, there is growing evidence that an important link between PD and T2DM does exist (Fiory et al., 2019; Sharma et al., 2021). In fact, T2DM and PD share characteristic phenotypes such as mitochondrial dysfunction, increased OS levels, inflammation and ER stress, which eventually might lead to the activation of the apoptotic pathway (Hassan et al., 2020; Rocha et al., 2021). Moreover, brain is one of the major targets of insulin; therefore, dysregulation of the insulin signaling pathway (ISP) may exert detrimental effects in neurons, and even be the cause of cell death (Hong et al., 2020). In addition, due to the common phenotypes found between both diseases, several antidiabetic drugs are being tested as potential PD treatments (Hassan et al., 2020; Renaud et al., 2018; Sharma et al., 2021).

Animal models have become powerful tools to study and identify pathogenic mechanisms underlying human diseases like PD and DM (Lilao-Garzón et al., 2021; McGurk et al., 2015; Muñoz-Soriano & Paricio, 2011). In this scenario, we work with a *Drosophila* PD model based on inactivation of the *DJ-1β* gene (ortholog of human *DJ-1*). Previous studies demonstrated that *DJ-1β* mutant flies displayed PD-related phenotypes like motor defects and reduced lifespan. Moreover, these flies showed alterations in the activity of enzymes involved in the defense against OS and, in consequence, increased OS marker levels such as several reactive oxygen species (ROS) and protein carbonylation (Casani et al., 2013; Lavara-Culebras et al., 2010; Lavara-Culebras & Paricio, 2007). Furthermore, we have also shown that *DJ-1β* mutant flies display metabolic disturbances reflected by an increase in the glycolytic pathway and alterations in several metabolite levels (Solana-Manrique et al., 2020, 2022). *Drosophila* has also emerged as an excellent model for DM since it uses triacylglycerols and glycogen to store energy like humans. In addition, *Drosophila* synthesizes insulin-like peptides (ILP), which are secreted by specialized neurons, named insulin-producing cells, and glia in the brain (Álvarez-Rendón et al., 2018). There are seven ILPs that are partially redundant (Álvarez-Rendón et al., 2018; Pasco & Léopold, 2012); however, ILP2, 3 and 5 are the most important considering glycaemia control (Gáliková & Klepsatel, 2018). Moreover, *Drosophila* regulates glucose homeostasis through the ISP, which is evolutionary conserved and contains similar elements and regulatory interactions than those found in human ISP (Teleman et al., 2012). To date, several strategies have been developed to model T2DM in this organism (Liguori et al., 2021; Morris et al., 2012). One involves feeding flies with an hypercaloric medium, in which sugar, protein or lipid content is increased with respect to regular food; alternatively T2DM can be modeled by silencing several genes like those encoding ISP components, among others (Liguori et al., 2021; Morris et al., 2012).

In this study, we have used PD and T2DM model flies, a mouse T2DM model and neuron-like human cells to identify common links between both diseases. The *Drosophila* T2DM model is a diet-induced one in which wild-type flies were fed with high sugared medium (Morris et al., 2012); the mouse T2DM model is a well-established one based on high-fat diet (Surwit et al., 1988), which is the best one reflecting the human disease as it develops together with obesity. While in flies, simple sugars from food are taken up passively from the digestive tract, in mice the excess of energy from the hypercaloric diet is stored as fat, inducing obesity. Fat accumulation leads to insulin resistance, hyperglycemia, and ultimately T2DM (Graham & Pick, 2017). Our results showed that *Drosophila* PD and T2DM models exhibit common phenotypes like alterations in carbohydrate metabolism, mitochondrial dysfunction or motor defects, among others. In addition, we demonstrated that both fly and mouse T2DM models presented DA neurodegeneration, a hallmark of PD. We have also confirmed that increased glucose levels have a detrimental effect in human neuron-like cells by triggering apoptosis, which leads to cell death. Besides, the hypercaloric medium used to generate T2DM model flies exacerbated those phenotypes in *DJ-1β* mutants, indicating that DJ-1 might have a role in carbohydrate homeostasis. Finally, we have confirmed that metformin, an antidiabetic drug, is a potential candidate for PD treatment and that it could prevent PD onset in T2DM model flies.

## MATERIAL AND METHODS

### Fly stocks and culture conditions

Fly stocks employed in this study were *y,w* (Bloomington Drosophila Stock Center #6598: *y*^*1*^,*w*^*1118*^) and the *DJ-1β*^*ex54*^ strain (referred to as *DJ-1β*) from the J. Chung laboratory (Park et al., 2005). Stocks and fly crosses were cultured using standard *Drosophila* medium at 25°C. Newly eclosed female flies were fed either with control medium (normal diet, ND) or with a high-sugar medium (high sugar diet, HSD), prepared using standard *Drosophila* food supplemented with sucrose to a final concentration of 30%. Flies were transferred to new vials every 2-3 days. When performing metformin treatments, vials with either control or high sugar medium contained a final concentration of 25 mM of this compound (Sigma-Aldrich).

### Rotenone exposure

For rotenone treatment, 1-day-old *y,w* female flies were cultured in standard *Drosophila* feed supplemented with 500 µM rotenone for 7 days (rotenone was dissolved in DMSO and a stock of 100 mM was prepared). Flies were transferred to new vials every 2 days.

### Determination of glycogen and soluble carbohydrates

Glycogen and soluble carbohydrates were calculated using a protocol adapted from (Foray et al., 2012). Briefly, groups of five 15-day-old female flies of each genotype and culture condition were homogenized in 200 µl of PBS with a steel bead using a TyssueLyser LT (Qiagen) for 2 min at 50 Hz. Fly extracts were further centrifuged at 180g for 10 min at 4 °C in order to discard debris. Next, 90 µl of the supernatant were collected, to which 10 µl of sodium sulphate 20% (w/v) and 750 µl of methanol were added. After vortexing the sample, it was centrifuged at 180g for 15 min at 4 °C. In this step, glycogen remained in the pellet and soluble carbohydrates in the supernatant. For glycogen estimation, pellet was washed twice with 80% methanol. Then, 1 ml of anthrone reagent (1.42 mg/ml in sulphuric acid 70% (v/v)) was added and samples were heated at 90 °C for 15 min. Subsequently, samples were cooled on ice and centrifuged. 200 µl of each replicate were added in a 96-well plate and absorbance was measured at 625 nm using an Infinite 200 PRO reader (Tecan). For soluble carbohydrates estimation, supernatant was evaporated until a volume of 20 µl remained, and 750 µl of anthrone reagent were next added. Samples were heated at 90 °C for 15 min and, subsequently, cooled on ice. 200 µl of each sample were transferred to a 96-well plate, and absorbance was measured at 625 nm. In all experiments, three replicates per each culture condition were carried out.

### Weight estimation

Groups of ten 15-day-old flies of each genotype and culture condition were weighed in a microbalance to estimate their total weight. At least, six replicates per each group of flies were studied.

### Measurement of ATP levels

ATP levels were measured as described in (Solana-Manrique et al., 2022) using the ATP Determination Kit (Invitrogen) following manufacturer’s instructions. Briefly, groups of five female flies were homogenized in 200 µl of reaction buffer (supplied by the commercial kit). Then, fly extracts were boiled 4 min and centrifuged at 18.500g for 10 min at 4 °C in order to discard debris. Subsequently, 5 µl of fly extracts were added to 100 µl of the standard reaction solution in a white 96-well plate and luminescence was measured using an Infinite 200 PRO reader (Tecan). All experiments were performed in triplicate.

### RT-qPCR analyses

Total RNA from ten 15-day-old T2DM model or control female flies was extracted and reverse transcribed as described in (Solana-Manrique et al., 2020). RT-qPCR were performed as in (Solana-Manrique et al., 2020), and the following pairs of primers were used: *tubulin* direct primer (5’-GATTACCGCCTCTCTGCGAT-3’); *tubulin* reverse primer (5’-ACCAGAGGGAAGTGAATACGTG-3’); *PI3K* direct primer (5′-ATTTGGACTACCTACGGGAA-3′); *PI3K* reverse primer (5′-TGCTTTTTCCTCCGTGTAG-3′); *InR* direct primer (5′-TTTCACGGAAGTCGAACATA-3′); *InR* reverse primer (5′-GACCTTAGCATAGCTCGG-3′); *chico* direct primer (5′-TATGCACAACACGATACTGAG-3′); *chico* reverse primer (5′-GACTCTGTTTTGGCTGACA-3′). *tubulin* levels were measured and used as an internal control for RNA amount in each sample. All experiments were performed in quadruplicate.

### Climbing Assays

Motor performance of flies was analyzed by carrying out a climbing assay as previously described in (Sanz et al., 2017). Briefly, groups of 10-20 female flies were transferred to graduated plastic tubes, acclimated for 1 min, gently tapped down to the tube bottom, and allowed to climb for 10 s. At least four groups of each condition were analyzed and the climbing ability was measured as the average height reached by each group after 10 s.

### T2DM mouse model

Mouse work was conducted in accordance with the animal ethics research committee (protocol OEBA-ULPGC 10/2019R1) from the University of Las Palmas de Gran Canaria. Brains from 10-month-old C57BL/6J female mice fed with either standard diet (ND) (Envigo, Global Diet 2014) or high fat diet (HFD) (60% energy from fat, D12492; Research Diets, New Brunswick, NJ) for 12 weeks (Surwit et al., 1988) were snap-frozen in liquid nitrogen and stored at -80 °C.

Brains were thawed and dissected to obtain the Caudate-Putamen region. 100 mg samples were homogenized in 1 ml of RIPA buffer supplemented with a protease inhibitor cocktail (PPC1010, Sigma-Aldrich) with an Ultra-Turrax T25 device. Then, lysates were sonicated (Bioruptor® Standard, Diagenode) on ice for 20 min in 30 s pulses and centrifuged 10.000 g for 20 min.

### Cell Culture and Metformin Treatment

*DJ-1*-deficient and pLKO.1 control SH-SY5Y cells previously generated by our laboratory (Sanz et al., 2017) were cultured in selective growth medium consisting of Dulbecco’s Modified Eagle Medium/Nutrient Mixture F-12 (DMEM/F-12) (Biowest) supplemented with 10% (v/v) fetal bovine serum (Capricorn), 1% non-essential amino acids, and 100 mg/ml penicillin/streptomycin (Labclinics) at 37 °C and 5% CO_2_. Cell viability after supplementation with different concentrations of glucose or MET was evaluated using an MTT (3-(4, 5-dimethylthiazol-2-yl)-2-5-diphenyltetrazolium bromide) assay, as previously described in (Solana-Manrique et al., 2020).

### Quantification of Protein Carbonyl Group Formation

Protein carbonylation levels were measured in 15-day-old flies cultured in ND and HSD containing 0.1 % DMSO for untreated control experiments or supplemented with a final concentration of 25 mM metformin for treatment experiments. Protein carbonyl groups were measured in fly extracts using 2,4-dinitrophenyl hydrazine derivatization in 96-well plates (Greiner 96-well plate, polypropylene) as previously described in (Solana-Manrique et al., 2020). All experiments were carried out using three biological replicates and three technical replicates for each sample.

### Western Blotting

Western blots were performed as described in (Solana-Manrique et al., 2020). Protein extraction of flies fed with either ND or HSD, and supplemented with either 25 mM MET or 0.1% DMSO as control, was adapted from (Solana-Manrique et al., 2020). Briefly, fifty 28-day-old fly heads were homogenized in 200 µl of 50 mM Tris–HCl, pH 7.4, with a steel bead in a TissueLyser LT (Qiagen) for 3 min at 50 Hz. Fly extracts were centrifuged at 14,500g for 10 min at 4 °C and supernatant was collected. Protein extraction of pLKO.1 SH-SY5Y cells treated with 125 µM of glucose or control medium was carried out as previously described in (Solana-Manrique et al., 2020). Protein extraction of mouse brain was carried out as commented above (see T2DM mouse model section). The primary antibodies used were anti-TH (1:1000, Sigma), anti-α-tubulin (1:5000; Hybridoma Bank; 12G10), anti-JNK, anti-phospho-JNK (Thr183/Tyr185) (1:1000, Cell Signaling) and β-actin (1:1000, Santa Cruz). Secondary antibodies used were anti-rabbit or anti-mouse HRP-conjugated (1:5000, Sigma). Quantifications of protein levels were performed with an ImageQuantTM LAS 4000mini Biomolecular Imager (GE Healthcare), and images were analyzed with ImageJ software (NIH).

### Enzymatic Assays

The enzymatic activities of phosphofructokinase (Pfk; EC 2.7.1.11), pyruvate kinase (Pk; EC 2.7.1.40), and hexokinase (Hk; EC 2.7.1.1) were measured using coupled enzymatic assays in extracts of 15-day-old flies fed with either ND or HSD, as previously described in (Solana-Manrique et al., 2020). All experiments were performed in triplicate.

### Statistical Analyses

The significance of differences between means was assessed using a t-test when two experimental groups were analyzed. In experiments in which more than two experimental groups were used, the statistical analysis was made using the ANOVA test and Tukey’s post-hoc test. Differences were considered significant when P < 0.05. Data are expressed as means ± standard deviation (s.d.).

## RESULTS

### *DJ-1β* mutant and rotenone-treated flies exhibit alterations in carbohydrate levels

Although the cause of PD is still unknown, several studies suggest that metabolic alterations might play an important role in developing the disease (Anandhan et al., 2017). Indeed, we have recently demonstrated that *DJ-1β* mutant flies displayed an increased glycolytic rate as well as changes in the levels of several metabolites when compared to controls (Solana-Manrique et al., 2020, 2022). Among them, we found an important increase in trehalose levels, which is the main circulating sugar in the *Drosophila* haemolymph (Solana-Manrique et al., 2022; Yasugi et al., 2017). To detect additional alterations in carbohydrate levels, we decided to quantify soluble carbohydrates and glycogen in our fPD model flies. Our results showed that 15-day-old *DJ-1β* mutants have increased levels of both (Fig. 1A-B). Interestingly, we found that 15-day-old *DJ-1β* mutants also exhibited reduced weight compared to control flies (Fig. 1C). Weight loss is often associated with an increase of energy expenditure, which could be explained by a switch from TCA cycle to glycolysis (Liesa & Shirihai, 2013; Wu et al., 2017). In fact, recent studies from our group confirmed this switch, since fPD model flies showed increased glycolytic rate and reduced activity of several enzymes of the TCA cycle (Solana-Manrique et al., 2020, 2022). Subsequently, we analyzed whether rotenone-treated flies, a well-established *Drosophila* sPD model (Coulom & Birman, 2004), showed similar phenotypes to *DJ-1β* mutants. Indeed, we found that these flies also exhibited changes in carbohydrate metabolism when compared to vehicle-treated flies (DMSO), reflected by increased soluble carbohydrates and glycogen levels (Fig. 1D-E). In addition, rotenone-treated flies showed reduced weight (Fig. 1F). Taken together, our results indicated that both fPD and sPD model flies displayed alterations in carbohydrate metabolism.

**Figure 1.**
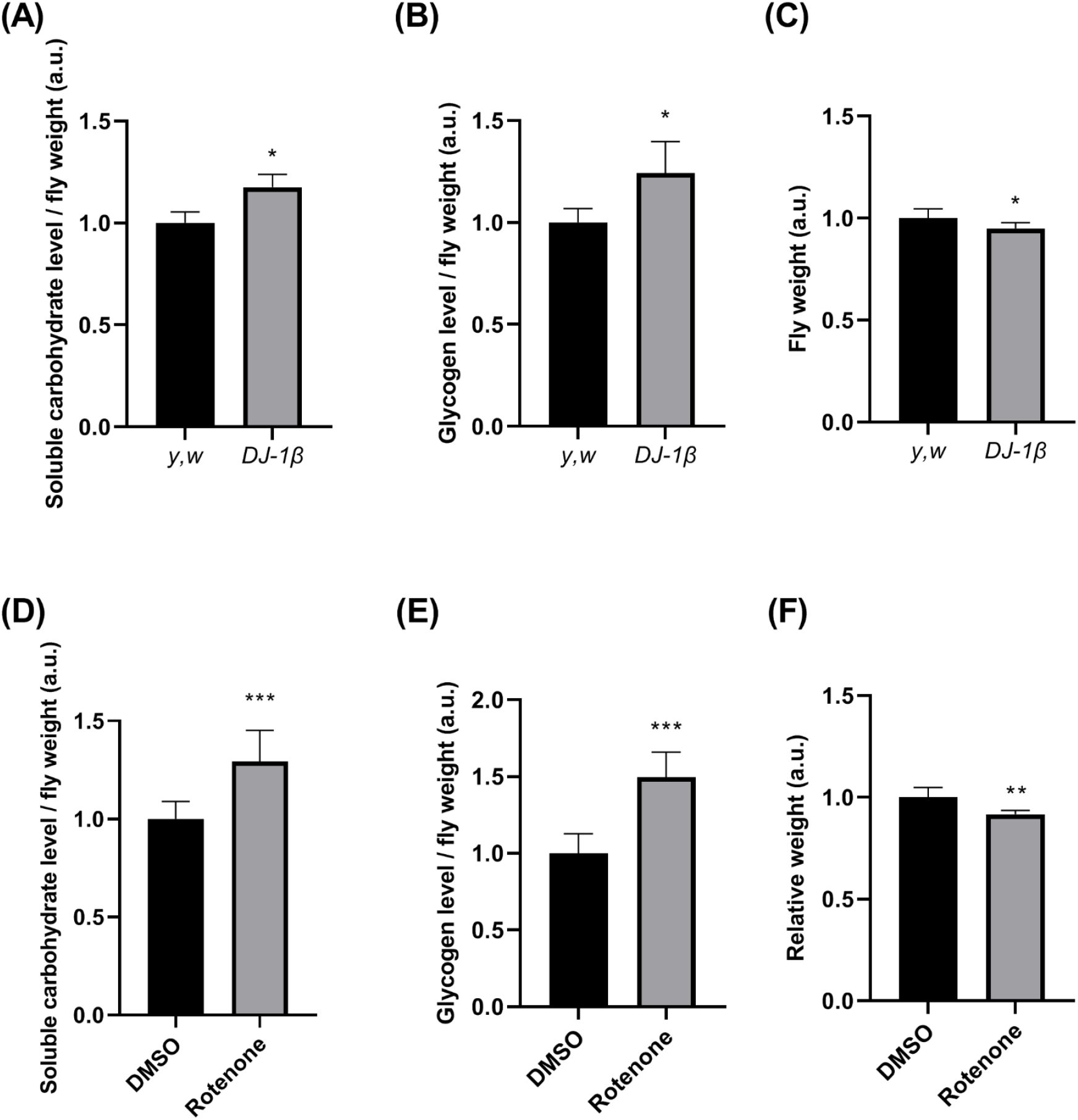
Metabolic alterations in PD model flies. **(A-B)** Soluble carbohydrate and glycogen levels in 15-day-old *DJ-1β*-mutant flies were analyzed using the anthrone method by absorbance. Fly weight of 15-day-old *DJ-1β* mutant flies. **(D-E)** Soluble carbohydrate and glycogen levels in 15-day-old rotenone-treated flies were analyzed using the anthrone method by absorbance. **(F)** Fly weight of 15-day-old rotenone-treated flies. Results were normalized to data obtained in *y,w* control flies in A, B and C; and to data obtained in flies cultured in control medium (DMSO) in D, E and F. Error bars show s.d. from three replicates and three independent experiments in A, B, D and E; and from at least six independent experiments in C and F (*, P < 0.05; **, P < 0.01; ***, P < 0.001).

### T2DM model flies display alterations in carbohydrate metabolism

High blood glucose level is a T2DM hallmark (Y. Zhang et al., 2021). Excitingly, several studies have revealed the existence of an association between T2DM and PD (Fiory et al., 2019; Sharma et al., 2021). While epidemiological links between both diseases have been deeply described, the underlying pathways and common mechanisms remain still unresolved (Cheong et al., 2020). In such a scenario, we decided to investigate the molecular and phenotypic links between T2DM and PD in *Drosophila*. Several *Drosophila* models for human T2DM have been described (Liguori et al., 2021). Among them, we used a diet-induced T2DM model in which wild-type flies were fed with a medium containing six times more sugar (HSD) than control medium (ND). A previous report showed that those flies developed insulin resistance, which is a hallmark of T2DM (Morris et al., 2012). To determine whether HSD-fed adult flies exhibited T2DM-related phenotypes we first measured carbohydrate levels. Our results indicated that 15-day-old T2DM model flies displayed increased soluble carbohydrates and glycogen levels (Fig. 2A-B). Subsequently, we found that these flies presented increased expression of *ILP2*, a gene encoding one of the three ILPs related to glycaemia control (Gáliková & Klepsatel, 2018) (Fig. 2C), thus confirming the existence of insulin resistance (Pasco & Léopold, 2012). In addition, we analyzed the expression of three genes encoding ISP components like *InR, PI3K* and *chico*. Our results showed that T2DM model flies exhibited increased the expression of those genes (Fig. 2D), indicative of pathway activity reduction (Pasco & Léopold, 2012). Taken together, increased levels of carbohydrates, the presence of insulin resistance, and decreased ISP activity validated flies fed with a HSD as a *Drosophila* T2DM model.

**Figure 2.**
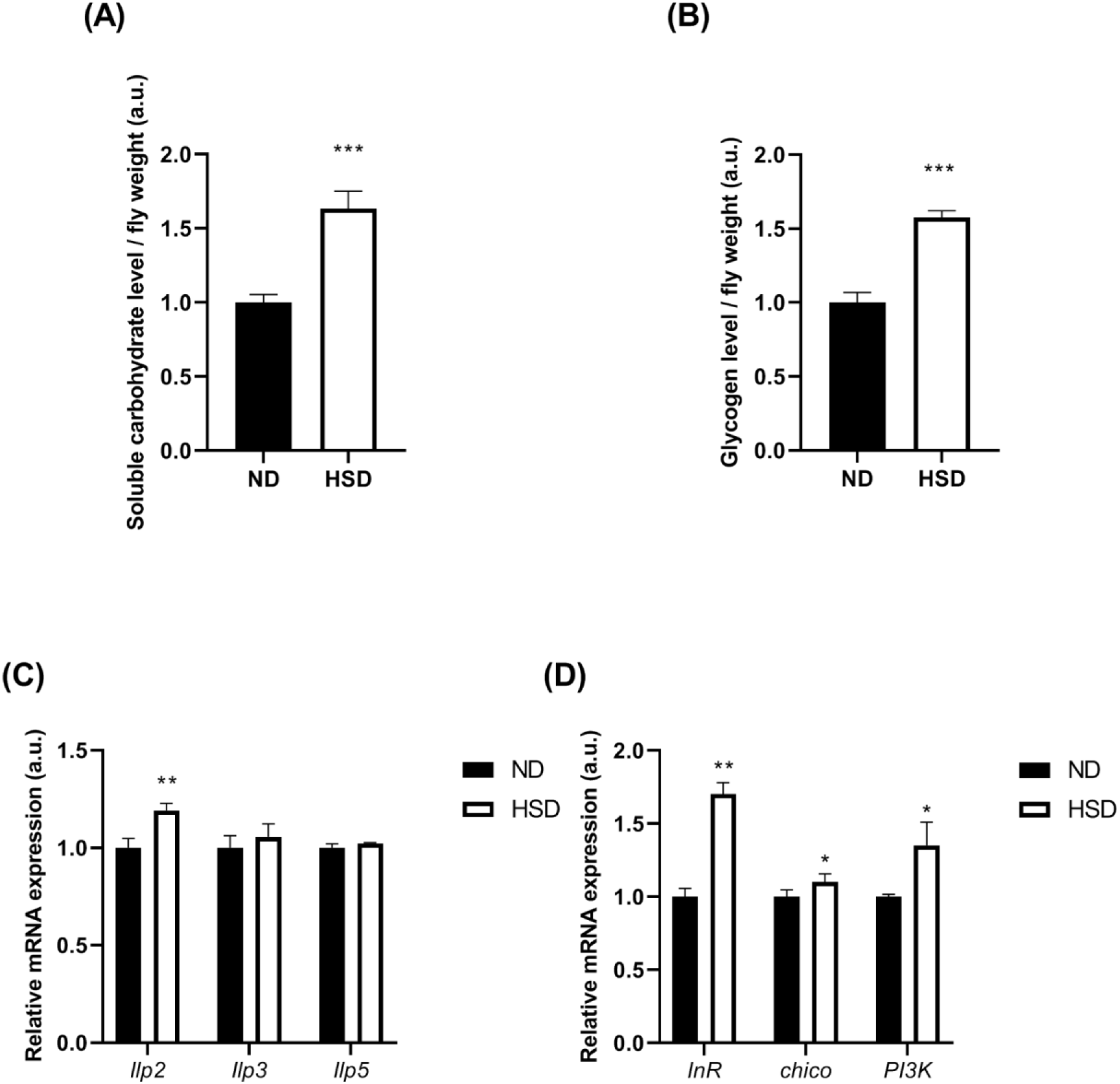
Metabolic alterations in T2DM model flies. **(A-B)** Soluble carbohydrate and glycogen levels in 15-day-old T2DM model flies (HSD) and flies fed with ND were analyzed using the anthrone method by absorbance. (**C-D)** Expression levels of *Ilp2, Ilp3* and *Ilp5* genes and of genes of the ISP in 15-day-old T2DM model flies were analyzed by RT-qPCR. In all cases, results are expressed as arbitrary units (a.u.) and are normalized to data obtained in flies cultured in ND. Error bars show s.d. from three replicates and three independent experiments in A and B, and from four independent experiments in C and D (**, P < 0.001; ***, P < 0.001).

### T2DM and PD model flies present common phenotypes

As indicated above, multiple studies highlight the link between PD and T2DM. Indeed, both diseases exhibit common phenotypes such as mitochondrial dysfunction (Hassan et al., 2020). To confirm this relation in our *Drosophila* models, we aimed to test whether T2DM model flies could present phenotypes similar to those observed in *DJ-1β* mutants. Mitochondria are the main ROS producers and their dysfunction leads to increased OS levels (Kudryavtseva et al., 2016). In fact, previous studies showed that *DJ-1β* mutant flies displayed elevated OS levels as well as alterations in antioxidant enzymes (Lavara-Culebras et al., 2010; Lavara-Culebras & Paricio, 2007). Hence, we decided to study OS levels in wild-type HSD-fed flies compared to controls (ND-fed flies). Our results demonstrated that 15-day-old T2DM model flies showed an increase in protein carbonylation levels (an OS marker) (Fig. 3A). In addition, it has been recently reported that mitochondrial alterations in *DJ-1β* mutant flies led to a reduction of ATP levels (Solana-Manrique et al., 2022). Consequently, glycolysis was enhanced in PD model flies in order to counteract the loss of energy production (Requejo-Aguilar et al., 2015; Solana-Manrique et al., 2020). Likewise, we have found that 15-day-old T2DM model flies presented a reduction of ATP levels (Fig. 3B), probably due to mitochondrial dysfunction. In addition, increased activity of hexokinase (Hk), phosphofructokinase (Pfk) and pyruvate kinase (Pk), enzymes involved in key regulatory steps of the glycolysis, were observed in these flies (Fig. 3C), suggesting an increased glycolytic rate. Moreover, we also found that T2DM flies presented reduced weight (Fig. 3D), likely due to an enhanced glycolytic activity as suggested for *DJ-1β* mutants.

**Figure 3.**
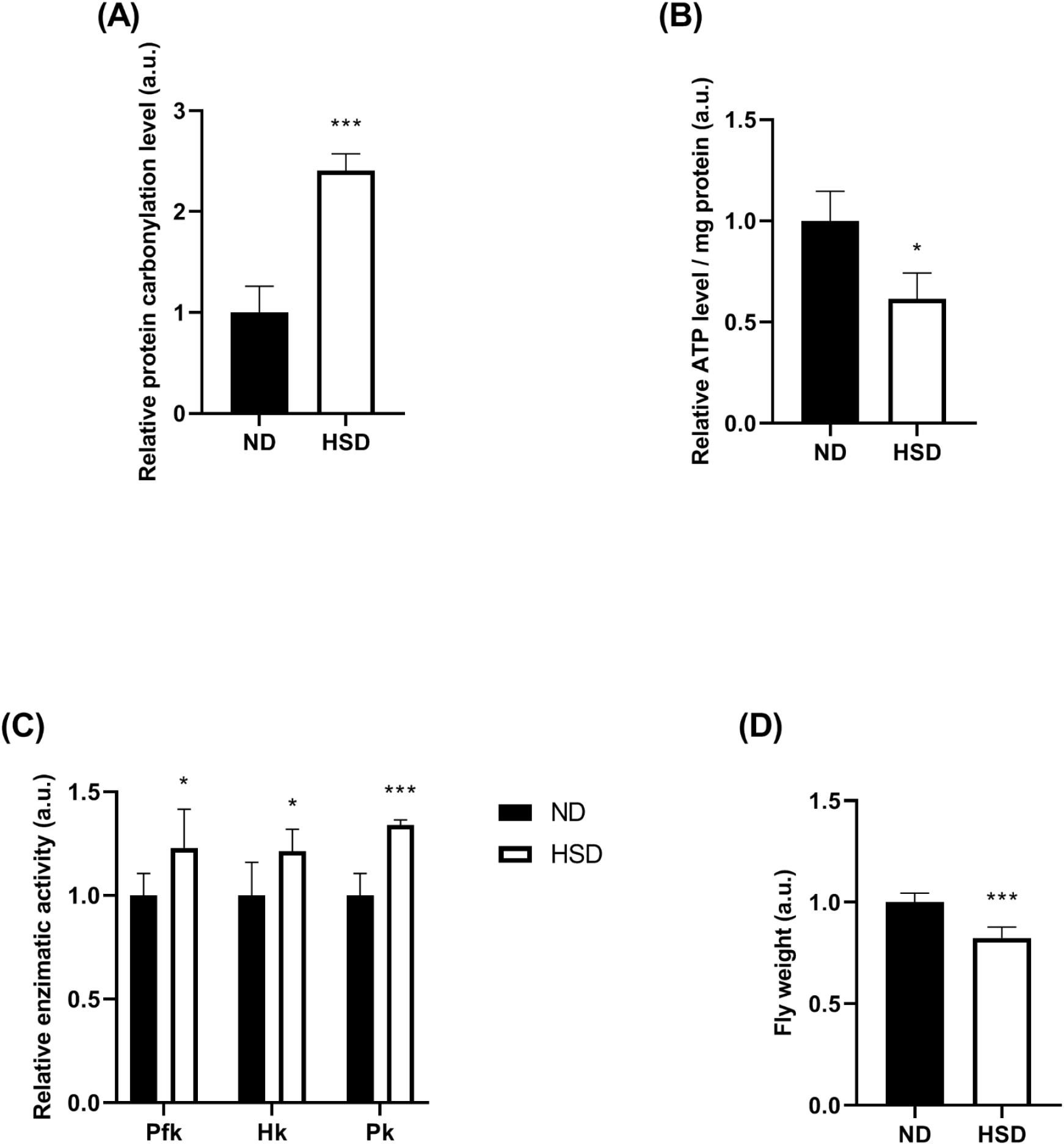
PD and T2DM phenotypes in T2DM model flies. **(A)** Protein carbonylation levels in T2DM model flies were analyzed by absorbance. Data were expressed as arbitrary units (a.u.) per mg of protein, and results were referred to data obtained in flies fed with ND (**B)** ATP levels in T2DM model flies were analyzed using the ATP Determination Kit (Invitrogen). (**C)** Activity of hexokinase (Hk), phosphofructokinase (Pfk) and pyruvate kinase (Pk) in T2DM model flies. Fly weight of T2DM model flies. Results were normalized to data obtained in flies fed with ND. Error bars show s.d. from three replicates and three independent experiments in A, B and C, and from at leats six independent experiments in D (*, P < 0.05; ***, P < 0.001).

On the other hand, T2DM is also considered as a risk factor of developing PD (Hassan et al., 2020). Among the most typical PD-related phenotypes exhibited by *DJ-1β* mutant flies, we found locomotor alterations and reduced life span (Lavara-Culebras et al., 2010; Lavara-Culebras & Paricio, 2007). In order to determine if T2DM might trigger PD onset, we decided to study these phenotypes in wild-type HSD-fed flies compared to controls (ND-fed flies). Our results demonstrated that 15-day-old T2DM model flies have a reduced life span (Fig. 4A). Besides, we found that flies fed with a HSD displayed locomotor deficits at 28 days of age, although no defects were found in younger flies (Fig. 4B). It was reported that motor symptoms appear in PD when there is approximately 50-60 % of DA neuron loss (Gallegos et al., 2015). Therefore, we decided to quantify levels of tyrosine hydroxylase (TH), a DA neurons marker, in T2DM model flies brain extracts to monitor DA neurodegeneration. For doing so, we performed western-blot analysis with 28-day-old T2DM model flies heads with an anti-TH antibody. Previous studies reported that TH levels are related to DA neuron content in fly brains (Molina-Mateo et al., 2017). Our results showed that TH levels were, in fact, reduced in 28-day-old T2DM model flies (Fig. 4C and Fig. S1); hence, confirming DA neuron loss in these flies. Taken together, our results validated the link between PD and T2DM in *Drosophila* models and supported PD development in T2DM.

**Figure 4.**
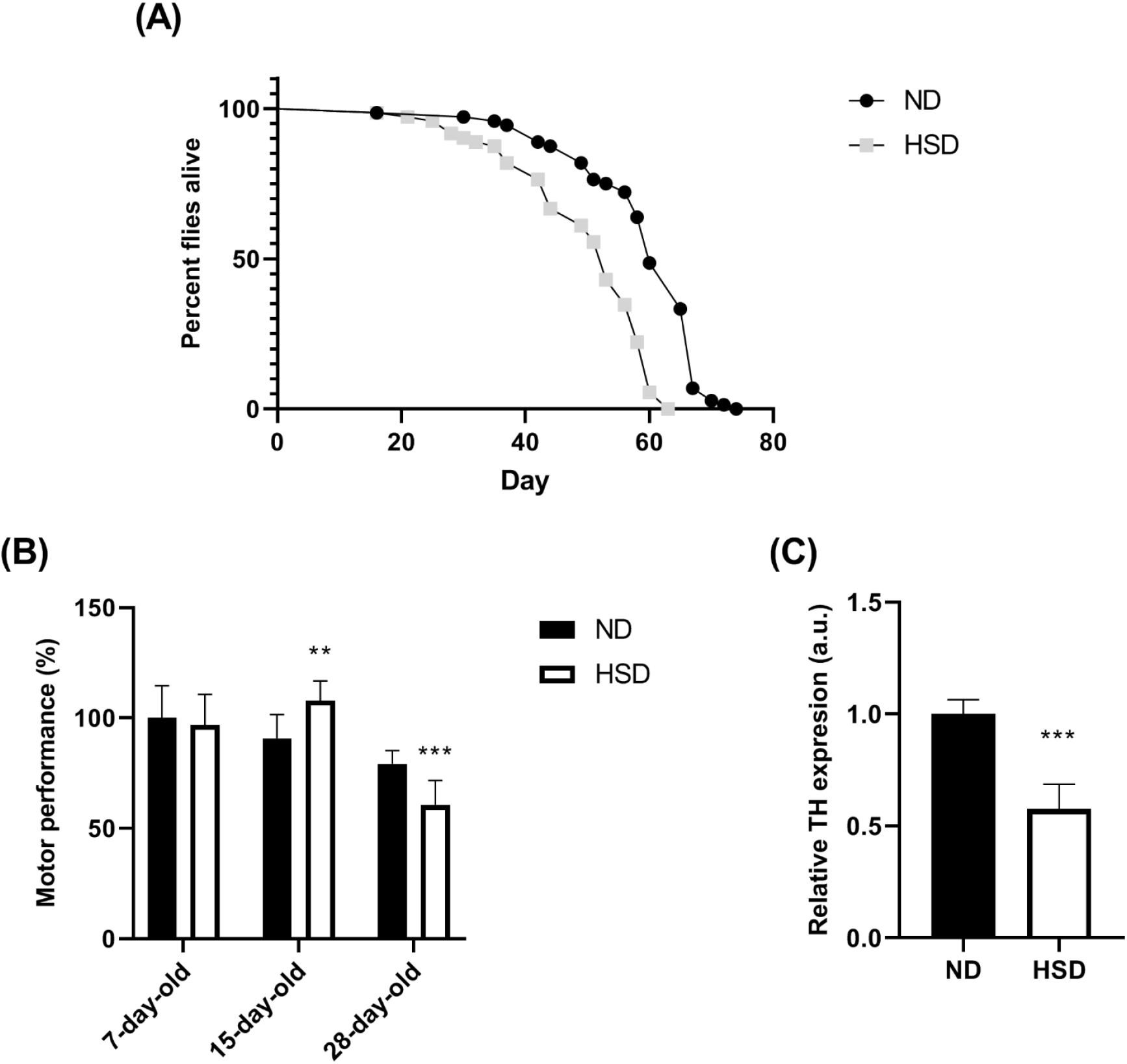
PD-related phenotypes in T2DM flies. **(A)** Comparison of survival curves of flies fed with HSD and ND. **(B)** Motor performance of flies fed with HSD and ND at different ages was evaluated performing a climbing assay, and results were normalized to data obtained in 7-day-old flies fed with ND. **(C)** Graphical representation of TH protein levels in T2DM flies. α-tubulin was used as loading control. Results are referred to data obtained in flies fed with in ND and are expressed as arbitrary units (a.u.). Error bars show s.d. from three replicates and three independent experiments (**, P < 0.01; ***, P < 0.001).

### *DJ-1*-deficiency leads to alterations in carbohydrate homeostasis

Several studies have demonstrated that modelling T2DM with HSD-fed animals resulted in increased OS levels, likely caused by high glucose concentration (Renaud et al., 2014). Interestingly, besides the already known antioxidant effect of the DJ-1 protein, there is growing evidence that it might also play an important role in metabolism (Mencke et al., 2021; Solana-Manrique et al., 2022). Therefore, it is plausible that loss of *DJ-1* function could be detrimental under hyperglycemic conditions. To confirm this, *DJ-1β* mutant flies were fed with a HSD during 15 days. Our results showed that they presented increased soluble carbohydrates as well as glycogen levels (Fig. 5A-B). To determine whether *DJ-1β* could play a significant role in their homeostasis, we studied if the increase of soluble carbohydrate and glycogen levels was affected by the HSD compared to ND (HSD/ND) in PD model and control flies. Our results showed that the increase of glycogen levels in HSD/ND was higher in *DJ-1β* mutant flies than in controls, while no differences were found in the increase of soluble carbohydrates (Fig. 5C-D). This suggests that *DJ-1β* may play an important role in glycogen metabolism. It has been reported that hyperglycemia increases OS levels (Renaud et al., 2014). Since *DJ-1β* mutants were shown to be hypersensitive to OS conditions (induced by paraquat, rotenone, H_2_O_2_, etc.) (Lavara-Culebras & Paricio, 2007; Meulener et al., 2005; Park et al., 2005), we analyzed if feeding PD model flies with a HSD had detrimental effects on motor ability. Accordingly, *DJ-1β* mutant flies cultured in those conditions displayed motor defects from day 7 (Fig. 5E), while controls fed with the same diet showed reduced motor ability at day 28 (Fig. 4B). These results confirm that *DJ-1β* mutant flies are more sensitive to increased carbohydrate levels, supporting an important role of *DJ-1β* in carbohydrate metabolism.

**Figure 5.**
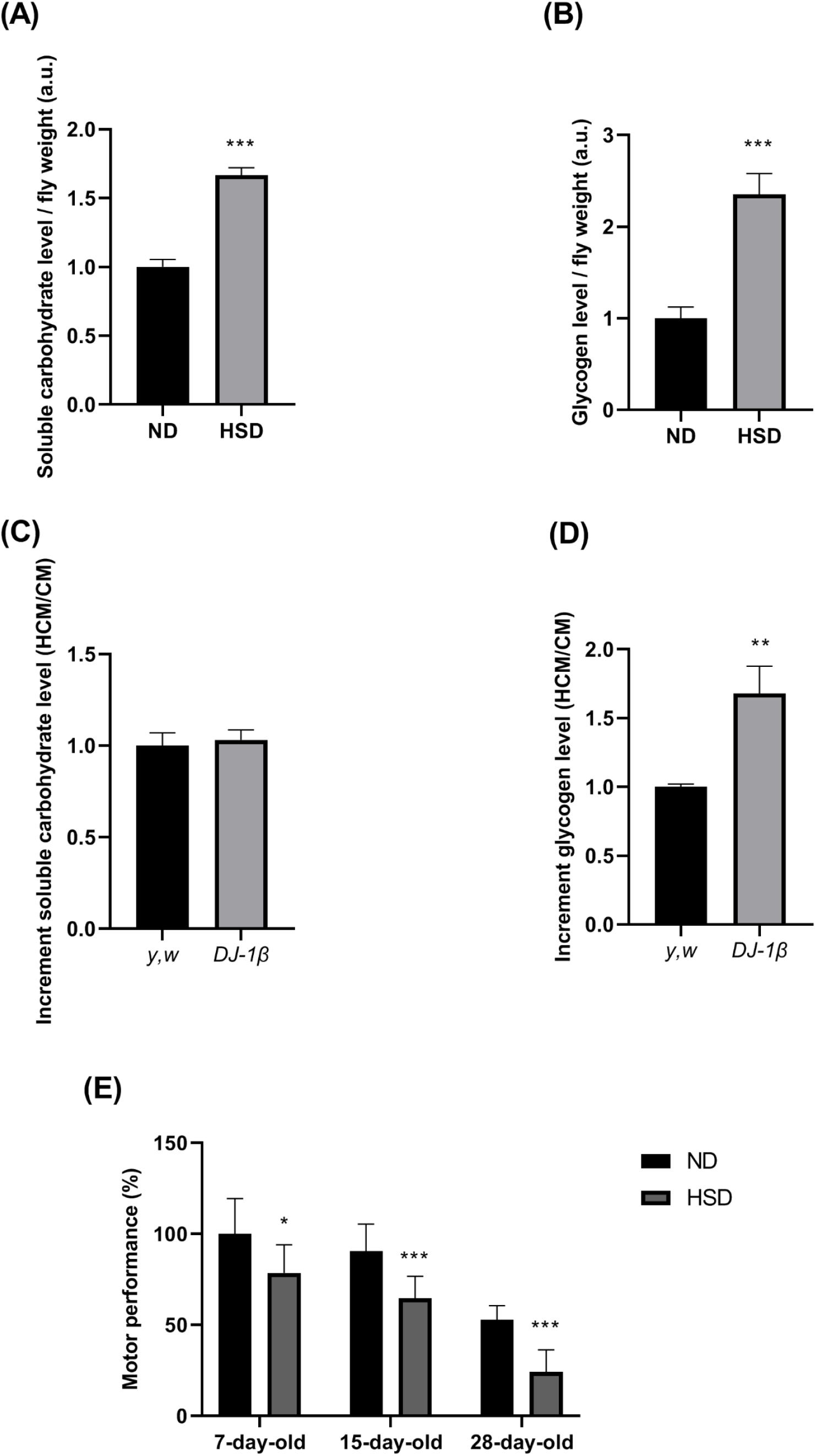
Effect of a HSD in PD model flies. **(A-B)** Soluble carbohydrate and glycogen levels in *DJ-1β* mutant flies fed with HSD and ND were analyzed using the anthrone method by absorbance. Results are expressed as arbitrary units (a.u.) per fly weight. **(C-D)** Increment of soluble carbohydrates and glycogen levels of *y,w* (control) and *DJ-1β*-mutant flies when fed with HSD compared to those flies fed with ND. **(E)** Motor performance of *DJ-1β* mutant flies fed with HSD and ND at different ages was evaluated performing a climbing assay and results were normalized to data obtained in 7-day-old flies fed with ND. In all cases, error bars show s.d. from three independent experiments in which three biological replicates were used (*P < 0.05; **P<0.01; ***P < 0.01).

### High glucose levels trigger neurodegeneration in mice and SH-SY5Y human cells

To further study the relevance of T2DM on PD development, we decided to use a T2DM mouse model. We used a well-stablished mouse T2DM model induced by 12 weeks of HFD which leads to glucose levels increase in serum (Surwit et al., 1988). As in the T2DM model flies, we found that 10-month-old female mice fed with HFD showed a TH expression reduction in brains when compared to ND-fed mice (Fig. 6 and Fig. S3). These results supported the onset of PD in these mice. Similar results have been obtained in other T2DM mouse models (Renaud et al., 2018).

**Figure 6.**
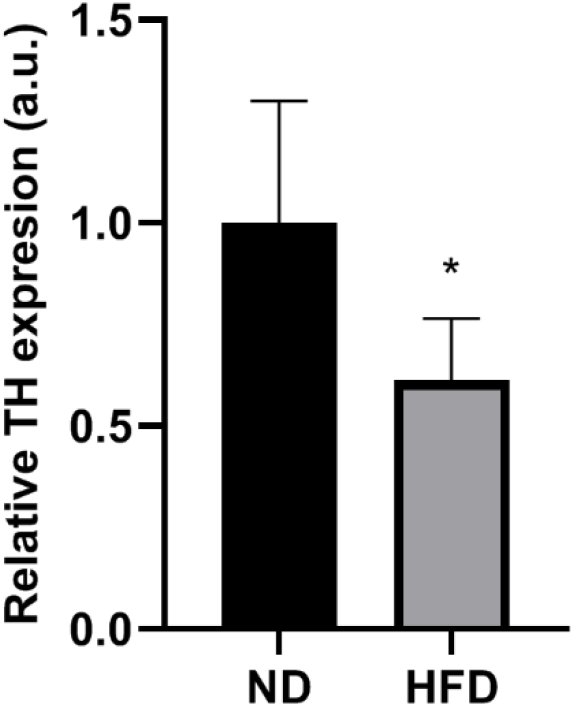
Tyrosine hydroxylase expression in T2DM mouse model brain. Graphical representation of TH protein levels in T2DM mouse brain. B-actin was used as loading control. Results are referred to data obtained in mice fed with ND and are expressed as arbitrary units (a.u.). Error bars show s.d. from five independent experiments (*, P < 0.05).

To date, the cause by which T2DM might be triggering PD development remains unclear. Vascular defects, increased levels of methylglyoxal, OS or glucose were described as potential causes (Hassan et al., 2020; Murillo-Maldonado et al., 2011; Renaud et al., 2018). Since antidiabetic drugs (compounds able to reduce glucose levels) are being found as potential PD treatments (Renaud et al., 2018), we aimed to study if elevated glucose levels might be detrimental for neuron-like SH-SY5Y cells. To do this, we treated these cells with increased glucose concentrations in a range of 50-175 mM (Cho et al., 2019) and subsequently performed MTT viability assays. Our results indicated that viability was significantly reduced 100 mM glucose onwards (Fig. 7A), hence confirming the toxic effect of high glucose levels in these cells. Subsequently, to study the molecular mechanism by which glucose might be producing cell death, we decided to measure the activation of the pro-apoptotic factor JNK (Ambacher et al., 2012; Sanz et al., 2021; Zhang et al., 2019) after glucose treatment. We found that cells treated with 125 mM glucose showed increased JNK phosphorylation levels when compared to cells treated with vehicle (Fig. 7B and Fig. S3), which ultimately leads to cell death. Together, results obtained in the T2DM mouse model and in neuron-like SH-SY5Y humans cells supported the relevance of high sugar levels in neuronal degeneration.

**Figure 7.**
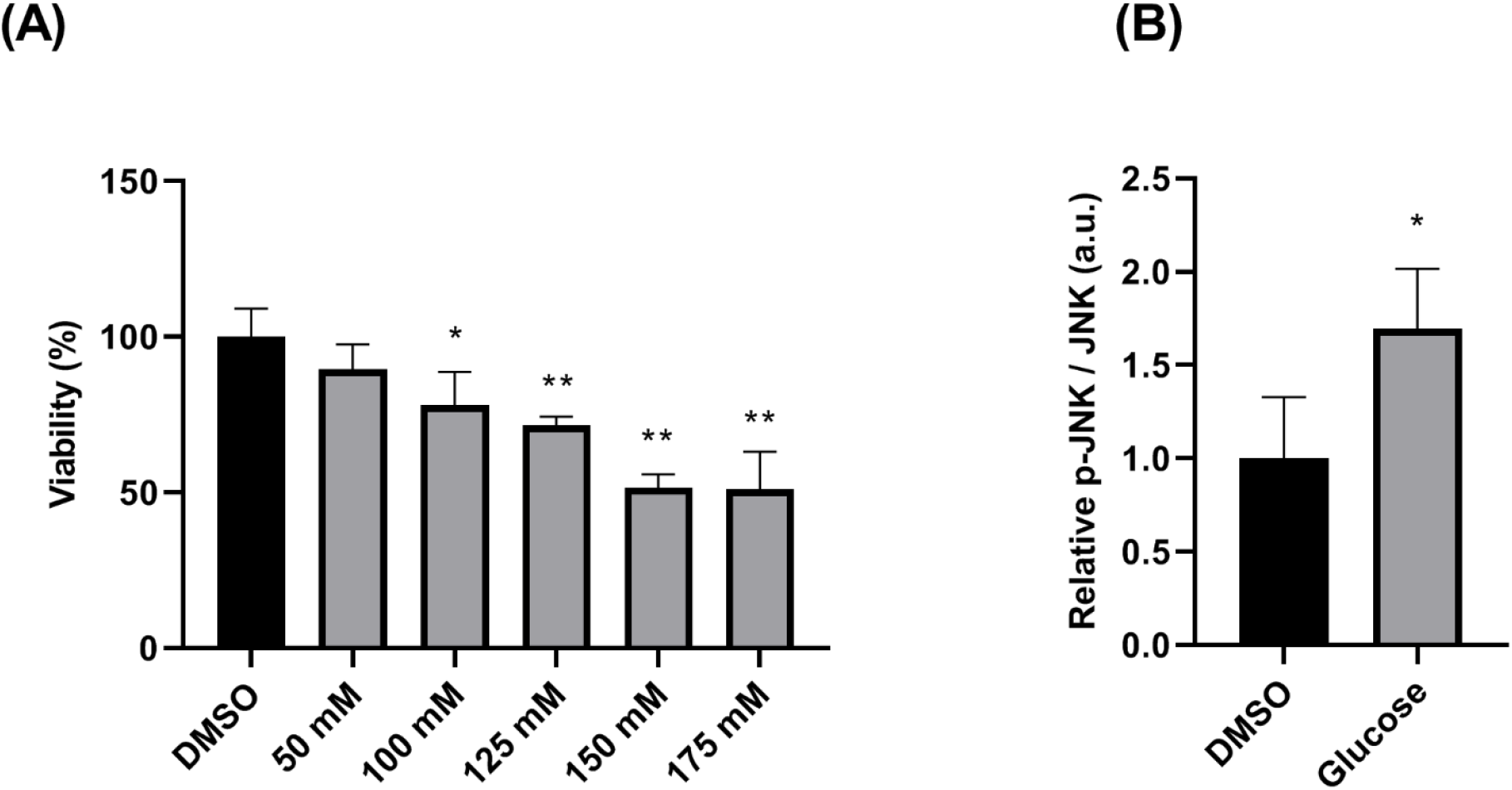
Effect of glucose on viability and JNK pathway activity in SH-SY5Y neuroblastoma cells. **(A)** Cell viability was measured in SH-SY5Y neuroblastoma cells treated with different concentrations of glucose (50-175 mM) or vehicle (DMSO), using an MTT assay. Results were normalized to data obtained in vehicle-treated cells (DMSO). **(B)** Antibodies against JNK, and p-JNK were used to detect proteins of interest in SH-SY5Y cells treated with vehicle medium (DMSO) or 125 mM glucose by Western blot. The relative ratio of p-JNK/JNK was analyzed by densitometry. Results are referred to data obtained in vehicle-treated cells and expressed as arbitrary units (a.u.). Error bars show s.d. from three independent experiments in which three biological replicates were used in A and from four biological replicates in B (*P < 0.05; **P < 0.01).

### Metformin ameliorates phenotypes in *DJ-1*-deficient human cells and T2DM model flies

Given the growing evidence of the relationship between PD and T2DM, antidiabetic drugs are being considered as promising PD therapies (Hassan et al., 2020; Renaud et al., 2018; Sharma et al., 2021). Interestingly, a pilot screen carried out by our group in *DJ-1β* mutant flies identified metformin as a candidate therapeutic compound for PD, being able to suppress motor defects in PD model flies (Sanz et al., 2017). Metformin (MET) is a well-known antidiabetic drug. However, it has been also reported to exert a neuroprotective effect through the 5′-adenosine mono-phosphate-activated protein kinase (AMPK) signaling pathway (Paudel et al., 2020). Therefore, we aimed to confirm this compound efficiency as PD therapeutic. First, we decided to determine whether MET was able to affect OS-induced lethality in our cell PD model based on *DJ-1* deficiency (Sanz et al., 2017). We found that MET reduced OS-induced cell death in *DJ-1*-deficient SH-SY5Y cells in a dose-dependent manner, being 100 µM the most effective concentration (Fig. S4). Subsequently, we tested if this compound was able to suppress DM as well as PD phenotypes in T2DM model flies. We found that HSD-fed flies treated with MET presented reduced levels of soluble carbohydrates and glycogen when compared to flies treated with DMSO as vehicle (Fig. 8A-B), hence confirming its antidiabetic activity. Subsequently, we also demonstrated that MET was able to suppress typical PD phenotypes, by reducing protein carbonylation levels (Fig. 8C), and improving locomotor activity of these flies (Fig. 8D). Accordingly, MET also increased TH expression in T2DM model flies (Fig. 8E and Fig. S1), which correlates with a reduction of DA neurodegeneration. These results confirm that MET might be a promising candidate therapeutic for PD.

**Figure 8.**
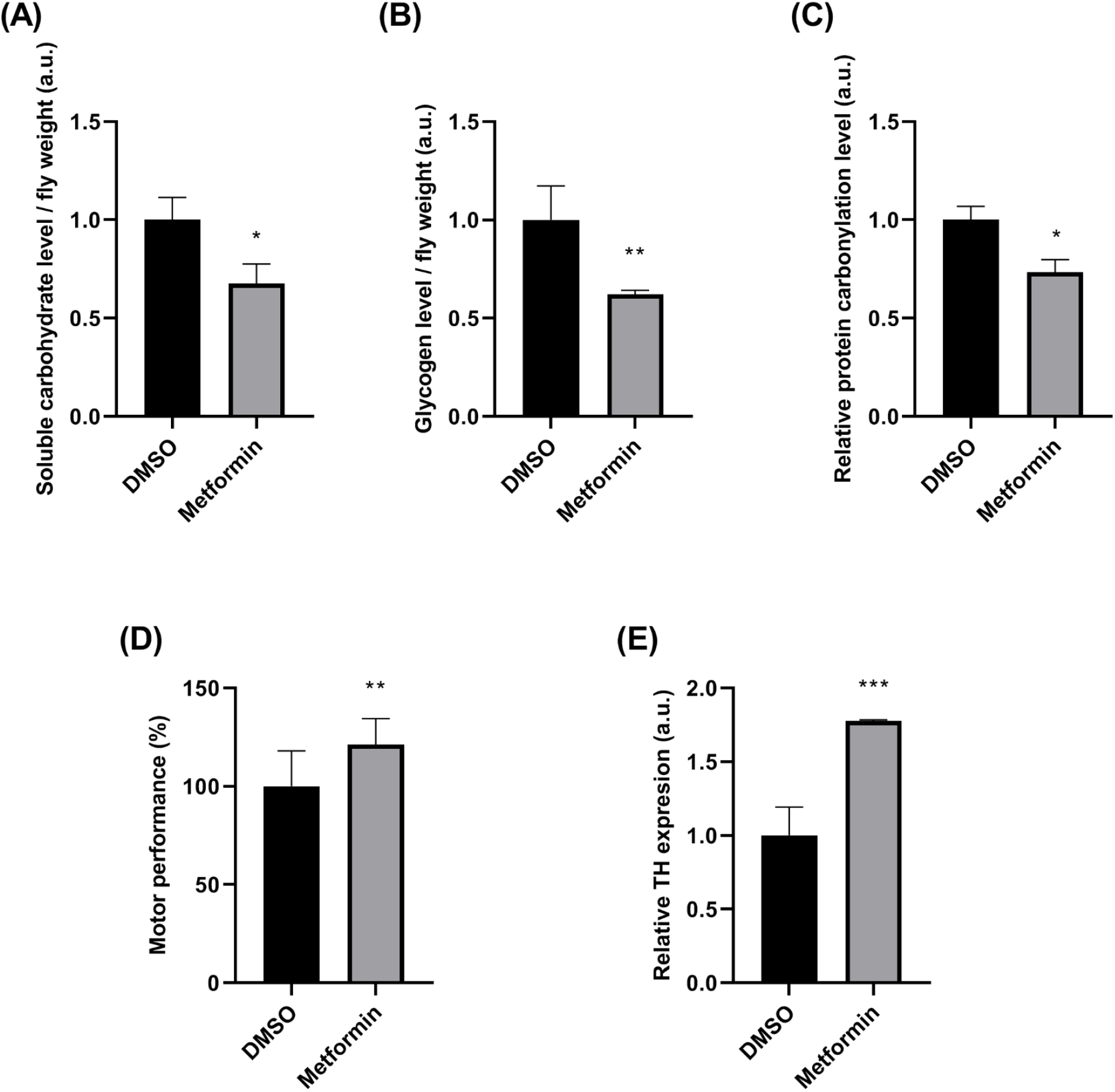
Effect of metformin in T2DM model flies. **(A-B)** Levels of soluble carbohydrates and glycogen in T2DM model flies treated with 25 mM MET were analyzed using the anthrone method by absorbance. Results are expressed as arbitrary units (a.u.) per fly weight and are normalized to data obtained in flies treated in vehicle medium (DMSO). **(C)** Protein carbonylation levels in T2DM model flies treated with 25 mM MET were analyzed by absorbance. Data were expressed as arbitrary units (a.u.) per mg of protein and results were referred to data obtained in T2DM flies treated with vehicle medium (DMSO). **(D)** Motor performance of T2DM model flies treated with 25 mM MET or DMSO was evaluated performing a climbing assay and results were normalized to data obtained in flies treated with vehicle medium (DMSO). **(E)** Tyrosine hydroxylase (TH) levels in T2DM model flies treated with 25 mM MET. Representative western blots are shown on upper panel. α-tubulin was used as control. Graphical representation of protein levels are shown on lower panel. Results are referred to data obtained in T2DM model flies treated with vehicle (DMSO) and are expressed as arbitrary units (a.u.). In all cases error bars show s.d. from three replicates and three independent experiments (*, P < 0.05; **, P < 0.01).

## DISCUSSION

The increase in life expectancy is leading to an increase in age-associated diseases, like PD and DM (Fiory et al., 2019), causing important social as well as economic burdens (Afentou et al., 2019; Standl et al., 2019). PD prevalence increases with age and affects 1-3% of the population over 60 years (Tysnes & Storstein, 2017). As for DM, it has a prevalence of 8.8 % in the group of population between 20-79 years in 2017 (Standl et al., 2019). In addition, an increase in the number of cases of both diseases is expected in the future (Afentou et al., 2019; Standl et al., 2019). Interestingly, recent epidemiological studies suggest that T2DM and PD are probably related (Fiory et al., 2019). In particular, most of them demonstrated that T2DM is a risk factor of developing PD, and could worsen cognition in PD patients (Hassan et al., 2020). Accordingly, an increase in ISP activity was reported to improve motor and cognitive abilities, and to exert neuroprotective effects in PD patients (Sharma et al., 2021). Besides, alterations found in chronic T2DM patients, like microvascular complications, may lead to nephropathy, retinopathy and neuropathy (Álvarez-Rendón et al., 2018). However, it was also proposed that neuropathy might be produced in T2DM by other mechanisms, such as hyperglycemia (Murillo-Maldonado et al., 2011). Regarding this, we recently showed that *DJ-1β* mutant flies presented higher levels of trehalose, the main circulating sugar in the *Drosophila* haemolymph (Solana-Manrique et al., 2022; Yasugi et al., 2017). Accordingly, we demonstrated in this work that these fPD model flies also present higher levels of soluble carbohydrates as well as glycogen, as also found in sPD model flies. Therefore, it is worth to consider that elevated carbohydrate levels might exert a detrimental role in developing PD.

In such a scenario, and to investigate the consequences of hyperglycemia and its relationship with PD, we used *Drosophila* as a model. *Drosophila* is a useful model to study T2DM due to the conservation of ISP between flies and humans (Teleman et al., 2012). In this work, we have used a diet-induced T2DM *Drosophila* model in which wild-type flies were fed with a HSD. T2DM model flies were reported to present higher levels of glucose than flies fed with ND, and to develop insulin resistance (Birse et al., 2010; Morris et al., 2012; Teleman et al., 2012). Accordingly, our results showed that T2DM model flies exhibited high levels of soluble carbohydrates as wells as glycogen. Additionally, we also confirmed that T2DM model flies showed a reduction of ISP activity and an increase of *ILP2* expression, which along with the increase of carbohydrate levels, are hallmarks of T2DM. Our results also showed that HSD-fed flies were smaller than those fed with ND, which could be explained by an increase of energy expenditure caused by the switch of metabolism from TCA to glycolysis (Liesa & Shirihai, 2013; Wu et al., 2017). This assumption was confirmed by finding increased activity of key enzymes of the glycolytic pathway in such flies. Moreover, mitochondrial defects have been reported in T2DM models (Hassan et al., 2020), which suggests a reduction in TCA activity. Excitingly, this work confirmed that ATP levels were also reduced in T2DM model flies. The finding of common phenotypes in T2DM model flies and *DJ-1β* mutants clearly supports the relationship between both diseases (Solana-Manrique et al., 2020, 2022).

Many studies have been recently performed to evaluate the link between PD and T2DM. In fact, most of the epidemiological studies conclude that there is an important risk of developing PD in T2DM patients, although there are also contradictory results (Fiory et al., 2019). It was also reported that hyperglycemia increases OS levels (Renaud et al., 2014), and promotes proteins glycation, which leads to their aggregation (Hassan et al., 2020). Moreover, it has been shown that insulin plays an important role in the central nervous system, since neurons use glucose as the main energy substrate (Hong et al., 2020; Renaud et al., 2018), acts as a pro-survival factor and regulates mitochondrial homeostasis (Fiory et al., 2019). Overall, these results support that T2DM is a risk factor of developing PD. Here, we have confirmed that T2DM model flies exhibited PD-related phenotypes like reduced lifespan, locomotor defects but also DA neurodegeneration (inferred by reduced TH levels), which is a pathological hallmark of PD. According to our results in *Drosophila*, we have also shown that DA neuron loss is evident in a T2DM mouse model. Interestingly, DA neurodegeneration as well as motor defects have been reported in other T2DM animal models (Figlewicz et al., 1996; Renaud et al., 2018); hence supporting our findings. Several mechanisms have been proposed to explain neurodegeneration like vascular defects, increased methylglyoxal levels, OS or increased glucose levels (Hassan et al., 2020; Murillo-Maldonado et al., 2011; Renaud et al., 2018). Here, we demonstrated that viability in SH-SY5Y neuroblastoma cell was impaired by high glucose levels (as previously reported in Cho et al., 2019), which triggered the apoptotic pathway by promoting JNK phosphorylation. Therefore, our results confirm that elevated glucose levels play a detrimental role in neuronal survival.

DJ-1 is a multifunctional protein, mainly acting as antioxidant. However, it also plays important roles in metabolism and mitochondrial homeostasis (Ariga et al., 2013; Mencke et al., 2021). In mammals, insulin is produced in pancreatic β-cells (Cao et al., 2014), which are hypersensitive to OS since they express reduced levels of antioxidant proteins (Jain et al., 2012). Besides, mitochondrial integrity and functionality is also important for glucose-stimulated insulin secretion (Jain et al., 2012). Interestingly, it was shown that *DJ-1* is upregulated under hyperglycemic conditions, and that DJ-1 levels are reduced in pancreatic islets of T2DM patients (Jain et al., 2012). Therefore, *DJ-1* might play an important role in maintaining pancreatic β-cells homeostasis. In *Drosophila*, insulin is produced and secreted by specialized neurons, known as insulin producing cells (Álvarez-Rendón et al., 2018). Therefore, it would be interesting to determine if those *Drosophila* cells present similar characteristics than human pancreatic β-cells, and if *DJ-1β* plays an important role in their homeostasis. Since hyperglycemic conditions might result in an increase of OS levels, PD models based on *DJ-1* deficiency could be more sensitive to those conditions. In fact, it was reported that *DJ-1*-deficient mice showed glucose intolerance and reduced β-cell area (Jain et al., 2012). In the present study, we have found that *DJ-1β* mutant flies fed with a HSD presented increased levels of soluble carbohydrates and glycogen as well as motor defects at an earlier age than those fed with ND. We have also shown that the increase of glycogen levels was higher in PD model than in control flies when fed with a HSD, thus suggesting that loss of *DJ-1* function might produce alterations in glycogen metabolism. In fact, it was shown that hyperglycemia enhances the expression of genes involved in glycogen synthesis, such as *glycogen synthase*, and that glycogen levels can be related to alterations in β-cell metabolism (Ashcroft et al., 2017).

Since T2DM and PD exhibit common phenotypes such as mitochondrial alterations, increased levels of OS markers, neuroinflammation as well as ER stress (Hassan et al., 2020; Rocha et al., 2021), compounds addressed to restore these phenotypes might be useful for treating both diseases (Hassan et al., 2020). In fact, antidiabetic drugs such as PPAR (peroxisome proliferator-activated receptor) agonists, GLP-1 (glucagon-like peptide-1) receptor agonists and MET are being considered as promising drugs for treating PD (Hassan et al., 2020; Sharma et al., 2021). In addition, there are several clinical trials to repurpose antidiabetic drugs in PD patients (Renaud et al., 2018). Besides, a recent study indicated that T2DM patients treated with antidiabetic drugs showed reduced PD incidence (Fiory et al., 2019). Excitingly, a pilot screening assay carried out by our group identified MET as a compound able to improve motor performance in *DJ-1β* mutant flies (Sanz et al., 2017). Furthermore, it has been shown that MET also prevents cell death in another mouse model for T2DM (Lu et al., 2016). In this work, we have validated MET as a candidate compound for PD by confirming its effectivity suppressing OS-induced cell death in *DJ-1*-deficient cells. Besides, we have shown that MET is also able to suppress DM and PD-related phenotypes in T2DM flies. First, we have demonstrated that 15-day-old T2DM flies treated with MET showed reduced levels of soluble carbohydrates as wells as glycogen, hence confirming its antidiabetic effect. Subsequently, we have shown that MET is able to reduce protein carbonylation in 15-day-old T2DM model flies, to improve motor performance and to reduce DA neurodegeneration in 28-day-old T2DM flies.

In summary, we have used *DJ-1β* mutant flies, well-established *Drosophila* and mouse T2DM models and neuron-like human cells to investigate the link between PD and T2DM. Our results demonstrated that fly models share common phenotypes such as carbohydrate levels alterations, dysfunction in metabolic pathways related to energy production, and increased OS levels. In addition, we have shown that T2DM model flies develop PD-related symptoms with age, and that that hyperglycemia might be one of the causes that triggers DA neurodegeneration in T2DM, both in fly and mammalian models. Interestingly, our results showed that MET, an antidiabetic drug, suppressed PD-related phenotypes in T2DM model flies. Besides, it also ameliorated OS-induced cell death in *DJ-1*-deficient human cells, thus supporting its use as a candidate PD therapeutic. Taken together, our results support the link between both diseases, and validate antidiabetic drugs as potential PD treatments. Finally, we have confirmed that *DJ-1β* mutant flies are more vulnerable to hyperglycemic conditions. Therefore, we propose that loss of *DJ-1* function could exert a detrimental effect in T2DM, being able to accelerate PD development in T2DM patients. Further experiments will be required to confirm this hypothesis.

## Abbreviations

DA: dopaminergic
DM: Diabetes mellitus
ER: endoplasmic reticulum
fPD: familial Parkinson’s disease
HFD: high fat diet
HSD: high sugar diet
ILP: insulin-like peptide
ISP: insulin signaling pathway
MET: metformin
ND: normal diet
OS: oxidative stress
PD: Parkinson’s disease
ROS: reactive oxygen species
sPD: sporadic Parkinson’s disease
T1DM: Type 1 Diabetes mellitus
T2DM: Type 2 Diabetes mellitus
TH: tyrosine hydroxylase

## ACKNOWLEDGMENTS

We are grateful to Dr. Jongkyeong Chung and the Bloomington *Drosophila* Stock Center for providing fly stocks. Research at the NP lab is supported by the University of Valencia (Programa de Acciones Especiales de Investigación, UV-INV-AE-1553209). SMD is supported by the “Viera y Clavijo” Program from the Agencia Canaria de Investigación, Innovación y Sociedad de la Información (ACIISI) and the ULPGC and JLG by the ULPGC predoctoral program. Research at the SMD lab is supported by the ACIISI (CEI2019-02), Programa de Ayudas a la Investigación de la ULPGC, and ACIISI co-funded by FEDER Funds (ProID2020010013).

## FIGURE LEGENDS

**Figure S1. Tyrosine hydroxylase expression in T2DM model fly heads.** Representative western blot using anti-TH and anti-α-actin antibodies in heads of 28-day-old flies fed with ND and HSD, untreated or treated with 25 mM metformin.

**Figure S2. Tyrosine hydroxylase expression in T2DM model mouse brains.** Representative western blot using anti-TH and anti-β-tubulin antibodies in brains of 10-month-old mouse brains cultured in ND and HFD.

**Figure S2. JNK and p-JNK expression in SH-SY5Y neuroblastoma cells.** Representative western blot of using anti-p-JNK and anti-JNK antibodies in SH-SY5Y cells treated with vehicle (DMSO) or with 125 mM glucose.

**Figure S4. Effect of metformin in *DJ-1*-deficient SH-SY5Y neuroblastoma cells.** Cell viability was measured in *DJ-1*-deficient cells using the MTT assay in presence of OS (induced with 100 µM H_2_O_2_). Cells were treated either with vehicle (DMSO) or with MET (1-100 μM). Results were normalized to data obtained in vehicle-treated mutant cells (DMSO). In all cases error bars show s.d. from three replicates and three independent experiments (*, P < 0.05; **, P < 0.01).

