## Supplementary figures and images for "Exploring the link between Parkinson’s disease and Diabetes Mellitus in *Drosophila*"

### Supplementary figures

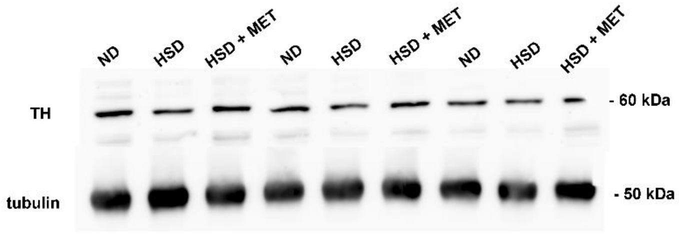

Figure S2

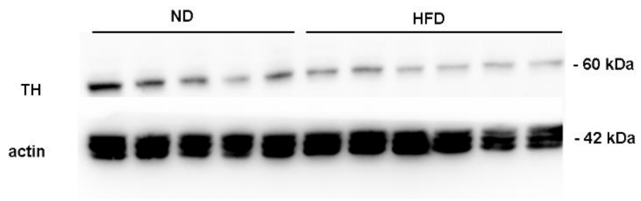

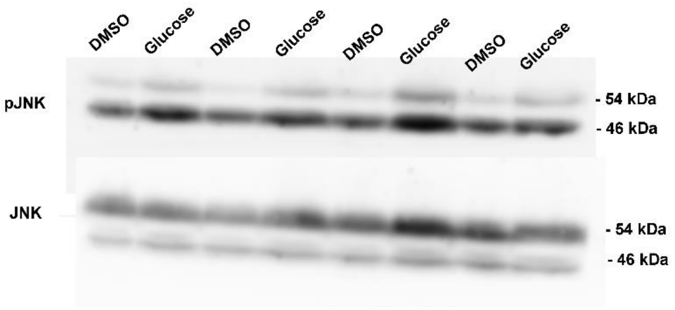

Figure S4

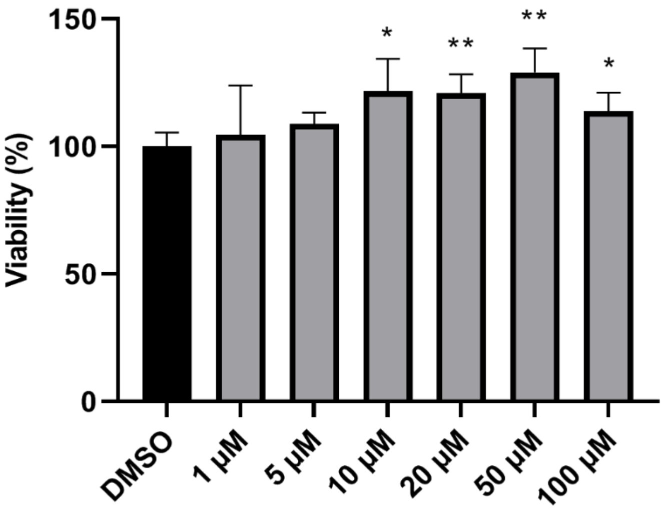
